# The architect of virus assembly: the portal protein complex nucleates procapsid assembly in bacteriophage P22

**DOI:** 10.1101/545707

**Authors:** Tina Motwani, Carolyn M. Teschke

## Abstract

The genetic material of tailed dsDNA bacteriophages, herpesviruses and adenoviruses is packaged into a precursor capsid through a 12-mer ring-shaped protein complex called the portal protein, located at a unique 5-fold vertex. In several phages and viruses, including T4, Φ29, and HSV-1, the dodecameric portal protein forms a nucleation complex with scaffolding proteins to initiate procapsid assembly, thereby ensuring incorporation of only one portal complex per capsid. However, for bacteriophage P22, the role of its portal protein in initiation of procapsid assembly is unclear. We recently developed an *in vitro* P22 assembly assay where portal protein is co-assembled into procapsid-like particles. We also showed that scaffolding protein catalyzes oligomerization of monomeric portal protein into 12-mer rings, and possibly forming a scaffolding-protein nucleation complex that results in one portal complex per P22 procapsid. Here, we present evidence substantiating that P22 portal protein, similar to the other dsDNA viruses, can act as an assembly nucleator. We find that the presence of P22 portal protein is able to increase the rate of particle assembly. Additionally, we show that P22 portal protein proper contributes to proper morphology of the assembled particles. Our results highlight a key function of portal protein as an assembly initiator, a feature likely conserved among these classes of dsDNA viruses.

## Importance

The existence of a single portal complex is crucial to the formation of mature infectious virions in the tailed dsDNA phages, herpesviruses and adenoviruses, and as such is a viable therapeutic target. How only one portal complex is selectively incorporated at a unique vertex is unclear. In many well-characterized dsDNA virus and phages, the portal acts as an assembly nucleator but early work by Bazinet and King (1) indicated that portal protein did not function this way in P22 procapsid assembly, leading to the suggestion that P22 might use a unique mechanism for portal incorporation. Here we show that portal protein does nucleate assembly of P22 procapsid-like particles. Addition of portal protein rings to an assembly reaction increases the rate of formation, the yield of particles, and corrects improper morphology. Our data suggest that procapsid assembly may universally initiate with a portal:scaffolding protein complex.

## Introduction

Double stranded DNA (dsDNA) tailed bacteriophages, herpesviruses, some dsDNA archaeal viruses and adenoviruses have a pseudo-icosahedral capsid with a portal protein complex incorporated at a unique five-fold symmetry axis (2–4). The portal complex is a ring of 12 identical subunits constituting an axial channel for the active packaging and ejection of the dsDNA viral genome (1, 5–8). The existence of a single portal complex is crucial to the formation of infectious virions. Evidence from dsDNA bacteriophages such as T4, SPP1, Φ29, and HSV-1 suggests that portal protein complexes play a key role in the nucleation and assembly of proper procapsids, thereby ensuring one portal per capsid (9–13).

The current hypothesized general mechanism for portal incorporation suggests that the portal complex interacts with the assembly chaperone, scaffolding protein. This scaffold-portal complex acts as a nucleus around which the coat protein monomers are polymerized, thereby ensuring that all procapsids have only one portal complex. This theory is supported by ample data from *in vivo* and *in vitro* studies of several bacteriophages. For instance, *in vivo* studies from bacteriophages have demonstrated interaction between scaffolding and portal proteins (12, 14–16). Additionally, *in vitro* studies in bacteriophage Φ29 has shown that the yield and rate of procapsid assembly increases in presence of portal protein, suggesting its role as an assembly nucleator (10). Similarly, data from HSV-1 indicates that portal complex gets integrated into the procapsids during initiation, thus highlighting its involvement of portal complex during early step of virus assembly (9).

Work of Bazinet and King (1988) indicated that bacteriophage P22’s portal protein might not nucleate P22 procapsid assembly because the presence or absence of portal protein did not affect the procapsid assembly kinetics *in vivo* (1, 17). Thus, P22 was suggested to employ a different mechanism to incorporate portal at the five-fold vertex, as compared to the other well-characterized dsDNA virus and phage systems. P22 is not unique in this observation as phage SPP1 also does not show a change in procapsid assembly kinetics in the absence of its portal, though for SPP1 portal protein does increase the yield of properly sized procapsids (12).

Recently we established an *in vitro* assembly assay for bacteriophage P22 where portal protein can be co-assembled into procapsid-like particles (PLPs) at a single vertex (18). Furthermore, we showed that scaffolding protein interacts with portal monomers and catalyzes the oligomerization of dodecameric portal rings. This interaction possibly leads to a formation of scaffolding-protein nucleation complex, that results in one portal complex per P22 procapsid.

In this report, we present evidence that P22 portal protein, in similar fashion to the portal proteins of other dsDNA viruses, can act as an assembly nucleator. As observed for bacteriophage Φ29 and HSV-1, we found that the presence of P22 portal protein is able to increase the rate of PLP assembly *in vitro*. Furthermore, the role of P22 portal protein as an assembly initiator was confirmed by its ability to fix the morphology of particles assembled at a high scaffolding protein to coat protein ratio. Our results highlight that the assembly initiator function of portal protein is well preserved among dsDNA viruses.

## Materials and Methods

### Purification of PC portal rings, SP and coat monomers for assembly reactions

Portal rings in the procapsid conformation (PC portal) were purified and assembled as described (18, 19). The portal protein concentration in all experiments is given as the 12-mer ring concentration. Scaffolding protein (SP) was purified as previously described (20). Coat protein monomers were prepared from empty procapsid shells. These shells were generated from an “assembler” plasmid containing only the genes for coat and scaffolding proteins, thereby eliminating the possibility of contaminating portal protein (21). The shells were denatured in 6.75 M urea, 20 mM HEPES pH 7.5 at 2 mg/mL final concentration for 30 min at room temperature (RT). The denatured coat protein monomers were diluted (1:1) with 20 mM HEPES pH 7.5, and extensively dialyzed against 20 mM HEPES pH 7.5 at 4 °C. Aggregates and uncontrolled shell assembly products were removed by ultracentrifugation at 221,121x g at 4 °C for 20 min in a Sorvall S120AT2 rotor.

### Procapsid-like particle assembly kinetics monitored by light scattering

For the assembly reactions, scaffolding protein was pre-incubated in a cuvette with PC portal rings for 5 minutes in 20 mM HEPES pH 7.5 with potassium acetate added for 70 mM buffer final concentration. The assembly reaction was initiated by the addition of coat protein monomers in 20 mM HEPES, pH 7.5 and 70 mM KAc buffer to a final concentration of 6.4 μM. The final scaffolding and portal protein concentrations were varied as described in the figure legends. The light scattering at 500 nm was monitored using a Horiba FluoroMAX 4 spectrofluorometer with the cuvette held at 20 °C. The excitation (Ex) and emission (Em) monochromators were set at 500 nm with slits set to 3.5 nm. A neutral density filter (OD 2.0) was placed in the excitation beam and the scattered light at 500 nm was recorded in counts per second (CPS) for 30 to 60 minutes.

### Agarose gel electrophoresis of *in vitro* assembled PLPs

The yield and oligomeric state of the PLPs and particles assembled with varied concentrations of scaffolding protein (3.0 to 44.7 μM) were analyzed by agarose gel electrophoresis. Twenty to thirty microliters of an *in vitro* assembly reaction were loaded onto 1.0% Seakem agarose gel in 1x TAE (40 mM Tris base, 20 mM acetate, 1 mM ethylenediamine tetraacetic acid (EDTA)). The gels were run at 100 V for ~75-90 minutes. The agarose gels were stained with Coomassie Blue and imaged with a Biorad Gel Doc Imager.

### Negative stain electron microscopy

Samples to be viewed by transmission electron microscopy (TEM) were prepared by applying 3 – 5 μL of *in vitro* assembled PLP (with or without PC portal rings) onto 300-mesh carbon coated copper grids (Electron Microscopy Sciences). Samples were absorbed for 1 minute and washed with 2-3 drops of water. Finally, they were stained with 1% uranyl acetate solution (wt/vol). The grids were visualized using FEI Technai G2 Spirit BioTWIN TEM equipped with an AMT 2k XR40 CCD camera at 68,000X magnification.

## Results

### Portal protein rings can nucleate procapsid assembly

For dsDNA bacteriophages such as T4, SPP1, Φ29, and HSV-1, portal protein complexes have a demonstrated role in the nucleation of procapsid assembly (9–13). On the other hand, *in vivo* studies in P22 showed that portal protein had no involvement in the initiation of procapsid assembly and no effect on the rate of procapsid assembly (1, 17), which led to the hypothesis that for P22 the portal protein is not involved in the nucleation of procapsid assembly. However, subsequent studies showed that excess portal protein *in vivo* led to aberrant structures as well as petite capsids, clearly indicating that portal protein affects morphology (17)

Having recently established conditions where we can incorporate portal rings into PLPs assembled *in vitro* (18), we decided to revisit the Bazinet and King (1988) result through *in vitro* assembly reactions. First, we tested the effect of portal rings on the kinetics of PLP assembly, as done previously for Φ29 portal protein (10). In all of the experiments, the procapsid conformation of portal rings (PC portal) was used (19). We pre-incubated scaffolding protein (9 μM) with varied concentrations of PC portal rings (0.0167 μM to 0.1 μM rings) in 20 mM HEPES pH 7.5 and 70 mM KAc buffer (final concentrations) in a cuvette. We kept the portal concentration low because of the effect of high concentration on morphology *in vivo* (17). The cuvette was incubated at room temperature (RT) for 5 minutes and the assembly was initiated by the addition of 6.4 μM coat protein monomers. The rate to PLP assembly in presence of PC portal rings was monitored by time dependent increase in the scattered light at 500 nm, measured in counts per second (CPS).

Both the rate of assembly and yield of PLPs increased with increasing portal protein up to 0.05 μM portal rings, after which portal decreased the rate of assembly (Figure 1). We determined the initial rate of each assembly reaction from the slope of the elongation phase directly after the initial lag phase as described (22), and plotted the rate relative to the rate of the assembly reaction conducted without added portal protein against portal concentration (Figure 1, inset). The positive effect of portal rings on the rate of capsid assembly suggests that bacteriophage P22 portal rings can act as an assembly initiator. High concentrations of PC portal rings (> 0.1 μM) decreased the yield of assembly (Figures 1). A similar result was observed when portal concentration was further raised to 0.2 μM rings (data not shown). We hypothesized that the decrease in the rate of PLP assembly at high concentration of PC portal rings (> 0.1 μM) could be due to interaction of portal protein with SP (18), thereby decreasing the amount of free SP for interaction with CP. Alternatively, too many nuclei could be forming and decreasing the concentration of coat protein available to assemble into PLPs. In the first instance, addition of extra scaffolding protein should relieve the inhibition of assembly, while in the second instance, we anticipated no effect on assembly by the addition of extra scaffolding protein.

**Figure 1.**
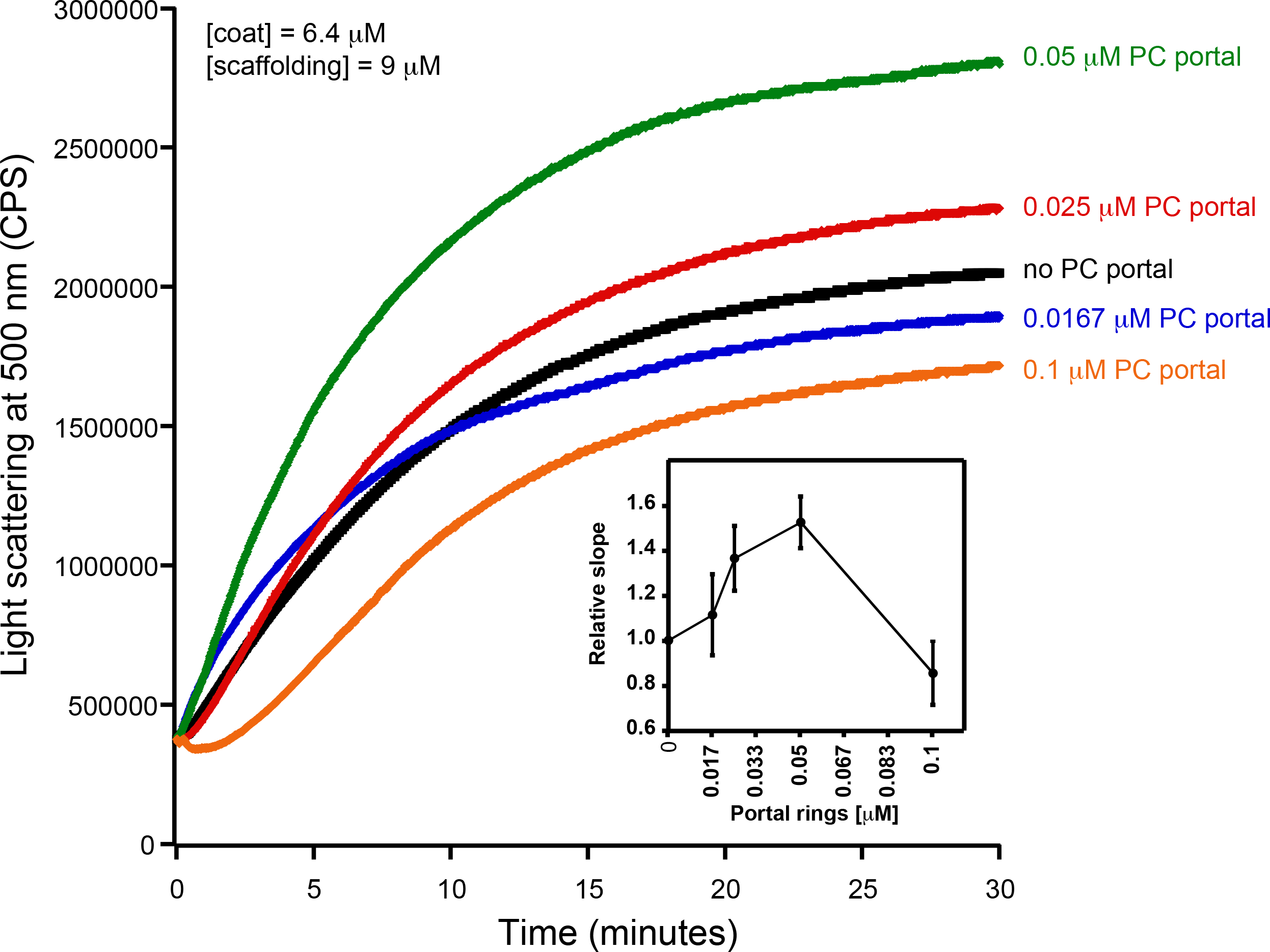
Portal protein rings can nucleate PLP assembly. PLP assembly kinetics in presence of varied concentration of PC portal rings was monitored by light scattering at 500 nm. The scaffolding protein (9 μM) and portal rings (0.0167 μM to 0.1 μM) were incubated in cuvette for 5 minutes in 20 mM HEPES pH 7.5 and 70mM KAc buffer. The assembly reaction was started by adding 6.4 μM coat protein monomers. The concentration of each protein used in the assembly reaction is indicated. The experiment was performed three independent times with different preparation of coat protein monomers. Shown are representative traces. **Inset:** Relative slope of the elongation phase directly after the initial lag plotted against portal concentration.

### Effect of high concentration scaffolding protein on PLP assembly

The above result (Figure 1) indicates that the appropriate concentrations of SP, CP and portal rings are critical for proper PLP assembly. In previous work, we showed that high concentrations of SP lead to aberrant structures *in vitro*, although these experiments were done in a different buffer than the HEPES Acetate buffer used here (23, 24). We have found that the HEPES acetate buffer used in the experiments here is so favorable for assembly that the critical concentration of coat protein is 0.075 mg/ml or 1.6 μM, which is four-fold lower that our previously characterized 20 mM phosphate buffer (26) (data not shown). Thus, we questioned if the positive effect of portal protein on assembly kinetics was due to an excess of scaffolding protein in the reactions, and that sequestration of some of the scaffolding protein by portal rings brings the assembly reaction back into balance. To test our idea, we performed a scaffolding protein titration experiment to determine the effect of varied scaffolding protein in PLP assembly, without portal protein added. We monitored the assembly of procapsids assembled at increasing scaffolding:coat protein ratios (SP:CP) with coat protein held at 6.4 μM by light scattering at 500 nm (Figure 2A). The relative slope of the early elongation phase of PLP assembly was plotted.

**Figure 2.**
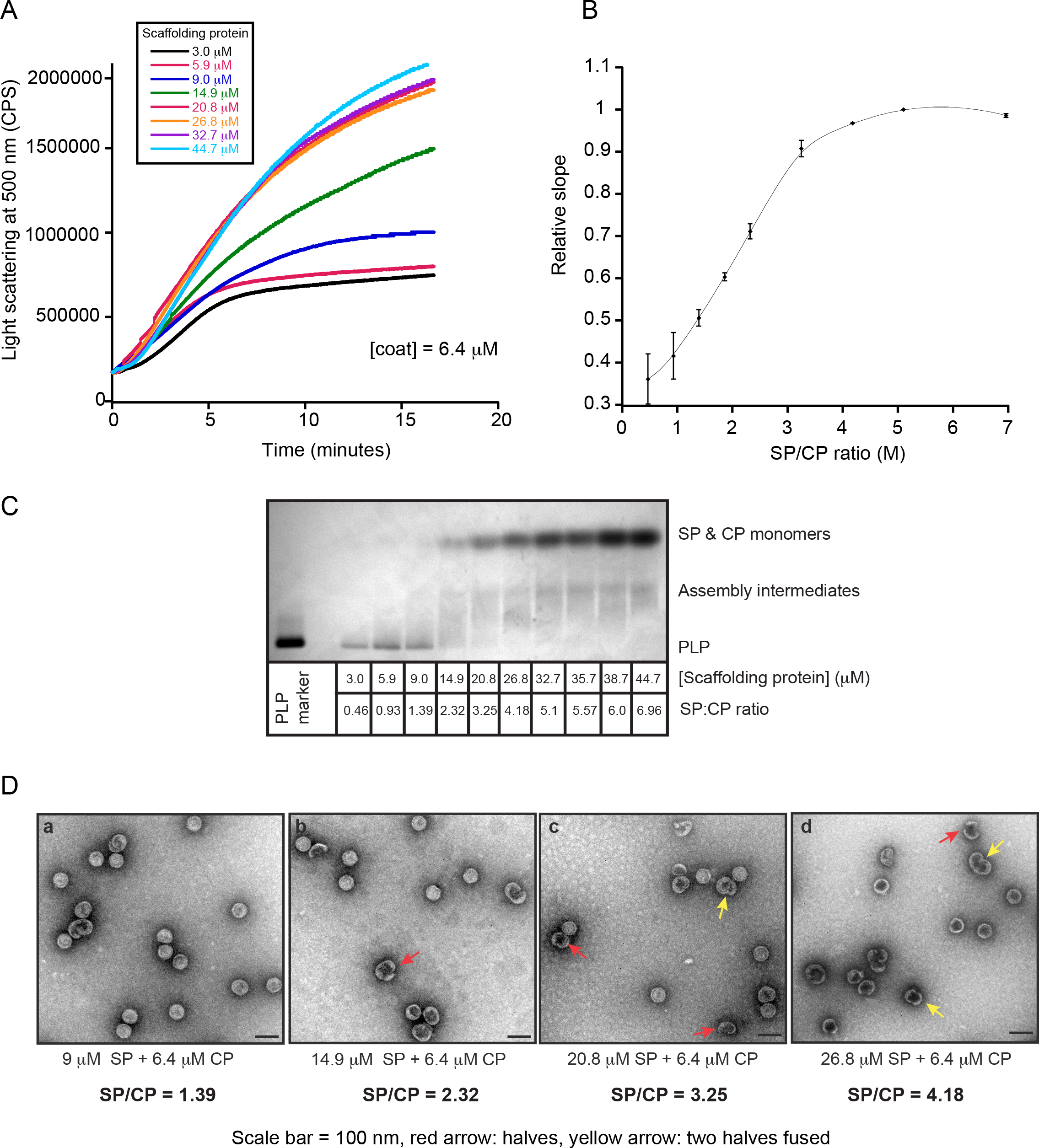
Effect of high concentration scaffolding protein on PLP assembly. (A) PLP assembly kinetics in presence of varied concentration of scaffolding protein monitored by light scattering at 500 nm. The scaffolding protein, from 3.0 μM to 44.7 μM, was incubated in cuvette for 5 minutes in 20 mM HEPES pH 7.5 and 70mM KAc buffer. The assembly reaction was started by adding 6.4 μM coat protein monomers. The experiment was performed three independent times with different preparation of scaffolding protein. Shown are representative traces. (B) Relative slope of the elongation phase directly after the initial lag plotted against SP/CP molar ratio. (C) PLPs assembled *in vitro* in the presence of varying concentration of scaffolding protein were run on a 1.0% agarose gel. The scaffolding protein concentration and the SP:CP ratio are indicated below the gel. (D) Negative-stain electron micrographs of *in vitro* assembled PLPs at varied SP:CP ratios. The scale bar is 100 nm. The red arrows indicate half-PLPs, and the yellow arrows indicate two half procapsids fused together.

Figure 2 shows that in our *in vitro* assembly conditions (20 mM HEPES and 70 mM potassium acetate buffer) that at low SP:CP ratios, scaffolding protein is limiting for assembly. Increased concentrations of SP yielded more particles until at higher scaffolding protein input concentrations, where SP is in large excess (>26.8 μM or SP:CP ratio of 4.18) the kinetics of assembly plateaued (Figure 2A and 2B). An aliquot of the assembly product obtained from each *in vitro* reaction was run on a native agarose gel, and the particles were visualized via transmission electron microscopy (TEM) (Figure 2C and 2D). At low SP:CP ratio (≤1.39), we observed a distinct PLP band on the gel and these particles had normal spherical procapsid-like morphology. However, at high SP:CP ratio (>1.39) the intensity of PLP band decreased and a band corresponding to assembly intermediates appeared. Examination of electron micrographs of these particles revealed the presence of bowl-shaped ‘half procapsid’ structures along with normal PLPs, as seen previously at a high SP:CP ratios (23, 24). These half procapsids increased in number with increasing SP concentration. In the experiments shown in Figure 1, the SP:CP ratio was 1.39. Addition of portal rings up to 0.05 μM increased the rate and yield of assembly. If the portal rings were sequestering scaffolding protein, based on the experiments shown in Figure 2, we should have observed a decrease in assembly rates because the reactions are in the regime where a change in available scaffolding protein would affect the rate (Figure 2B). Thus, our results show that there is a fine requisite balance between the critical protein components for assembly to occur properly.

### PC portal protein can act as an assembly nucleator even at high scaffolding and coat protein ratio

To further substantiate the role of P22 portal protein as an assembly nucleator, we investigated the effect of portal protein in assembly reactions when scaffolding protein is in excess, as described above. We incubated varied concentrations of PC portal rings (0.0267 μM to 0.2 μM) with scaffolding protein held at 20.8 μM (SP:CP ratio = 3.25) for 5 minutes in a cuvette at RT. We added coat protein monomers and observed the PLP assembly by light scattering. As shown in Figure 3, we found that even at this high SP:CP ratio, the presence of PC portal rings steadily increased the assembly kinetics. Maximal effect on assembly rate was observed with 0.053 μM or 0.075 μM PC portal rings in the reaction. Furthermore, we observed that at high concentration of PC portal rings (> 0.1 μM) the rate of PLP assembly gradually declined, similar to the effect noted when the scaffolding:coat protein ratio was low (Figure 1). In the high scaffolding concentration of this experiment, portal ring concentration (0.075 μM) is unlikely to bind enough scaffolding protein to limit PLP assembly. Together, our results (Figures 1 and 3) suggest that an optimal concentration of PC portal rings is required for its function as an assembly nucleator.

**Figure 3.**
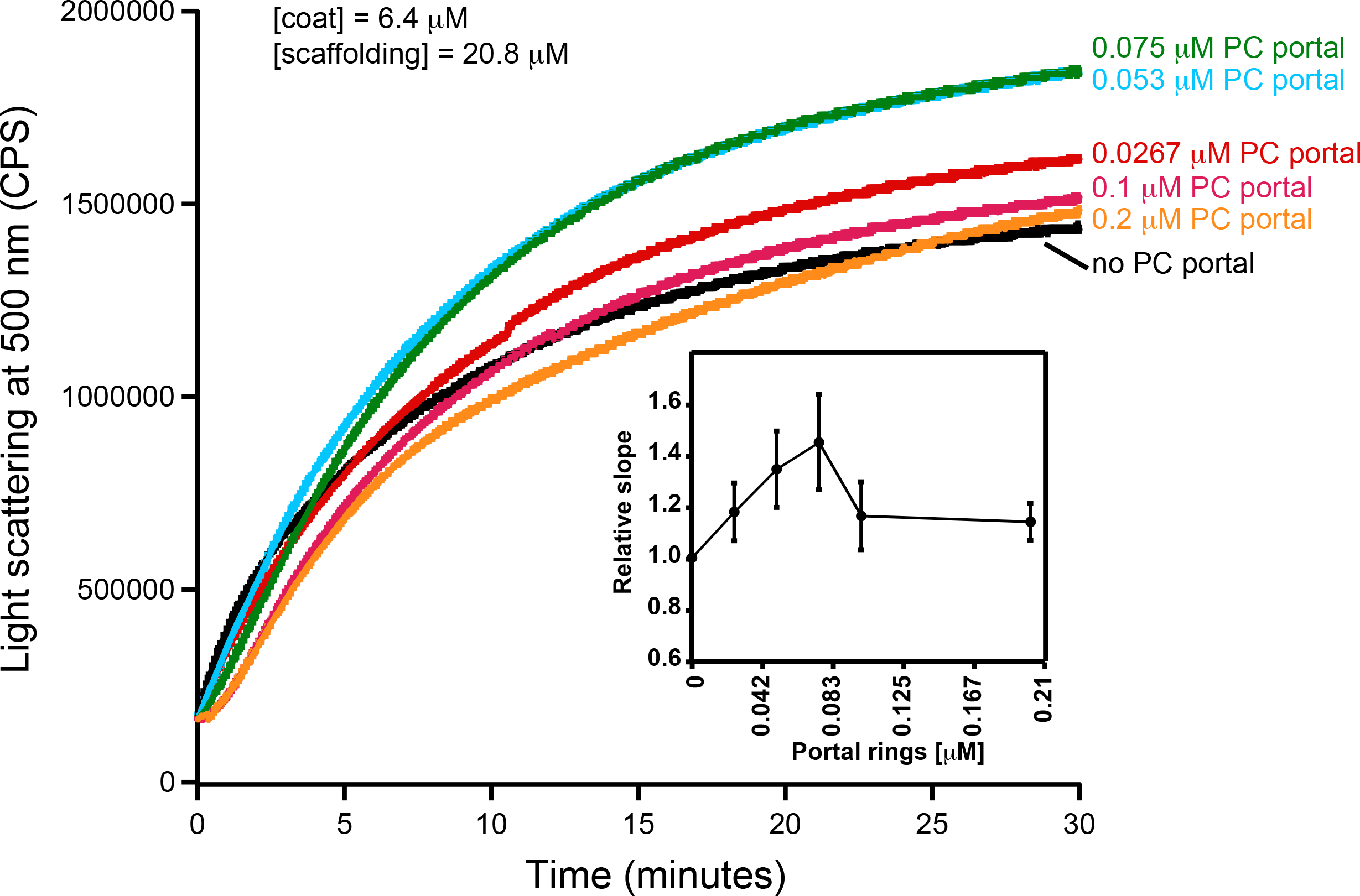
PC portal protein can act as an assembly nucleator even at high scaffolding and coat protein ratio. PLP assembly kinetics in presence of varied concentration of PC portal rings was monitored by light scattering at 500 nm. The scaffolding protein (20.8 μM) and PC portal rings (0.0 267 μM to 0.2 μM) were incubated in cuvette for 5 minutes in 20 mM HEPES pH 7.5 and 70mM KAc buffer. The assembly reaction was started by adding 6.4 μM coat protein monomers. The concentration of each protein used in the assembly reaction is indicated. The experiment was performed three independent times with different preparation of coat protein monomers. Shown are representative traces. **Inset:** Relative slope of the elongation phase directly after the initial lag plotted against portal concentration.

### Portal protein decreases aberrant structures

In other systems, portal protein can correct morphology defects in procapsid assembly (12, 16, 25–27). We tested if the presence of PC portal rings can complete the half-procapsids and aberrant structures observed at high SP:CP ratio (as shown in Figure 2). The partial PLPs are kinetically trapped structures, in this case because too many nuclei are forming so that coat protein becomes limiting and the particles are not able to complete (24, 28).

We hypothesize that the portal protein will affect the initiation of assembly by building a different nucleation complex with PC portal rings, scaffolding protein and coat protein monomers, thereby allowing more PLPs to form closed particles with a portal complex integrated at a single vertex, even at high SP:CP ratios. To test our theory, we incubated 0.059 μM or 0.1 μM PC portal rings with SP at 20.8 μM in 20 mM HEPES and 70 mM potassium acetate buffer for 5 minutes at RT (Figure 4A). As a mock control, we incubated only SP (20.8 μM) in our buffer alone. The assembly reaction was initiated by adding 6.4 μM coat protein monomers. This assembly reaction was allowed to reach completion and the PLPs purified over a Superose 6 gel filtration column. When AF488 labeled portal rings were used in the reactions, the labeled rings could be visualized in the PLP peak, as shown previously (Figure 4A) (18). The purified PLPs were analyzed by TEM prepared as described in Material and Methods. We analyzed over 2000 PLP particles from each reaction and quantified the number of complete and half/partial PLP structures in our micrographs. Consistent with our hypothesis, we observed the PLP particles assembled in presence of PC portal rings at high SP: CP ratio were more uniform and spherical as compared to without portal rings (Figure 4C). Additionally, the percentage of aberrant or half-procapsids was significantly reduced from 43% to 26% in presence of 0.059 μM PC portal rings and to 32% in presence of 0.01 μM PC portal rings, concomitant with an increase in normal sized and closed PLPs (Figure 4B). Thus, our data indicate, that similar to the other dsDNA viruses (HSV-1, Φ29, T4 and SPP1), bacteriophage P22 portal protein can act as an assembly initiator, a feature of portal protein that is likely conserved among all dsDNA viruses.

**Figure 4.**
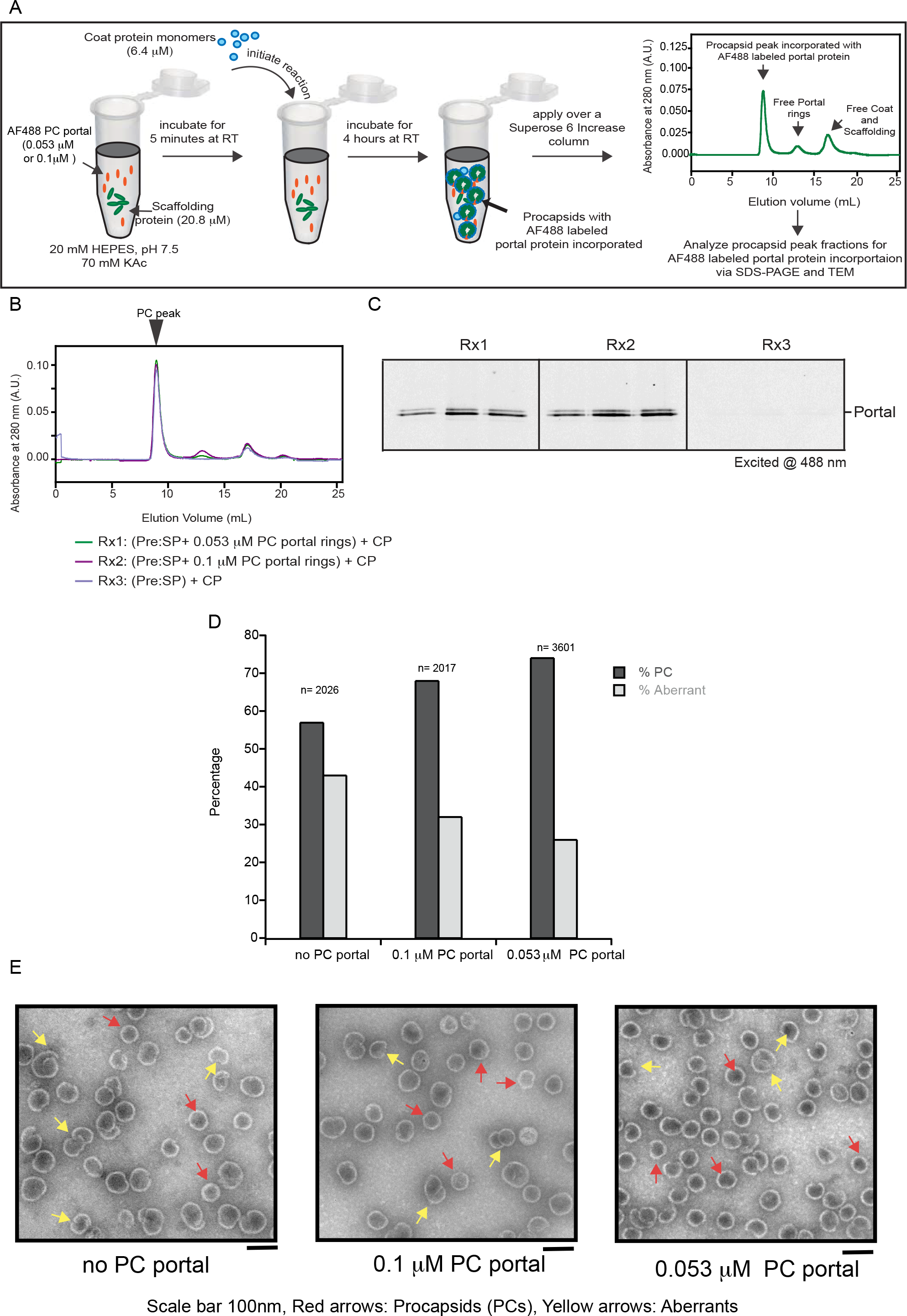
P22 portal protein completes half-procapsids assembled at a high SP: CP ratio. (A) Alexa Fluor labeled PC portal rings (0.059 μM or 0.2 μM) were incubated with scaffolding protein (20.8 μM) in 20 mM HEPES pH 7.5 and 70 mM KAc buffer for 5 minutes. CP monomers (6.4 μM) were added to initiate capsid assembly process. The reaction was allowed to reach completion at RT (4 hours) and the *in vitro* assembled PLP’s were separated by gel filtration chromatography. The purified peaks were analyzed by SDS-PAGE. (B) Superose 6 elution profiles of assembly reactions. Rx1: CP and SP with 0.0.059 μM AF488 labeled portal rings, Rx2: CP and SP with 0.1 μM AF488 labeled portal rings and Rx3: CP and SP alone (mock reaction). (C) SDS-PAGE gel of the purified PLP peak fractions from Rx1 to 3. The presence of labeled PC portal rings in these fractions was visualized using a Pharos FX plus Molecular Imager. (D) Percentage of uniform and spherical procapsids (black bars) and aberrant or half-procapsids (grey bars) assembled *in vitro* in absence (no PC portal) or presence of unlabeled PC portal rings (0.059 μM or 0.1 μM). More than 2000 particles were analyzed for each reaction. (E) Representative electron micrographs of the *in vitro* assembled PLPs assembled in absence or presence of unlabeled PC portal rings (0.059μM or 0.1 μM protomer). The red arrow indicates normal spherical PLP and yellow arrow indicates aberrant or half-procapsids. Scale bar, 100 nm

## Discussion

Portal proteins are essential to formation of infectious virions in the tailed dsDNA phages, herpesviruses and adenoviruses (2–4). The portal complex is the conduit for the DNA into the heads during packaging, and out of heads during infection. As such, portal protein is a potential therapeutic target.

The assembly of bacteriophage P22 is one of the most thoroughly studied of this class of viruses. Recently, we determined conditions that allowed the portal protein to be incorporated into assembling PLPs (18). This advance allowed us to revisit the role of portal protein in initiation of P22 assembly. Here, we show that the rate of assembly is increased by the addition of portal rings, as expected if portal rings are involved in initiation of PLPs (Figure 1 and Figure 3). We also demonstrated that the addition of portal rings could improve PLP morphology (Figure 4).

As with incorporation of HSV-1 and phage Φ29 portal complexes into PLPs, we found here that the protein concentrations need to be carefully balanced (9, 10). At low SP:CP ratios (<1.39) PLP assembly, with or without portal rings, occurs efficiently (Figure 5A). However, we observed that in the absence of portal rings at high SP:CP ratios (> 1.39), the PLP assembly is affected. The excess SP topples the balance of the critical components, resulting in rapid nucleation and formation of kinetically trapped metastable half-particles (Figure 5B). The presence of an optimal concentration of portal rings in assembly reactions carried out at high SP:CP ratio rebalances the reaction, resulting in in completion of the kinetically trapped incomplete shells (Figure 5B).

Over 30 years ago, Bazinet and King presented evidence that the absence portal protein did not affect the kinetics of P22 procapsid assembly *in vivo* (1). We believe this result could be due to: 1) a 3 min radioactive amino acid pulse that is nearly as long as PCs require to assemble *in vivo* (29, 30) that could obscure changed assembly rates; or 2) the sampling after the addition of chase was sparse so perhaps any changes in kinetics were simply not observed. We plan to revisit these *in vivo* experiments in the future.

**Figure 5.**
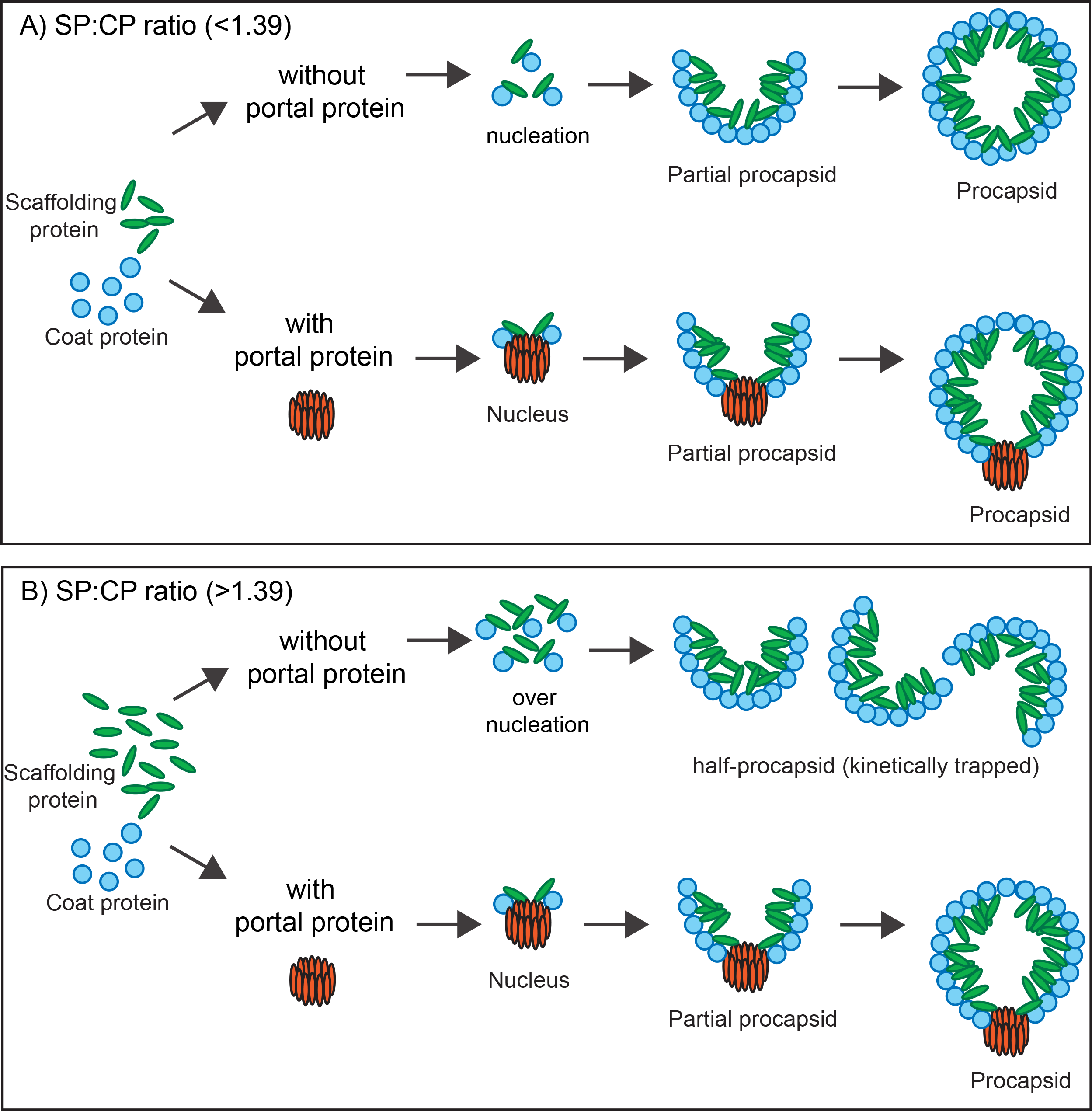
Cartoon showing the effect of PC portal rings on assembly at different SP:CP ratios.

## Acknowledgements

This work was supported by NIH grant GM07661 to CMT.

